## Supplementary for "Supramodal Shape Representation in the Human Brain"

**Supplementary Table 1. Demographic information of the early blind and their matched sighted control**

| Early Blind | Gender | Age | Handedness[a] | Onset of Blindness | Etiology | Sighted Control | Gender | Age | Handedness[a] |
|---|---|---|---|---|---|---|---|---|---|
| EB01 | F | 33 | 90 | Birth | Retinopathy of Prematurity | SC01 | F | 32 | 80 |
| EB02 | M | 32 | 60 | Birth | Retinitis Pigmentosa | SC02 | M | 29 | 50 |
| EB03 | M | 47 | 70 | Birth | Optic Nerve Hypoplasia | SC03 | M | 50 | 90 |
| EB04 | F | 31 | 70 | 8 Months | Retinitis Pigmentosa | SC04 | F | 31 | 40 |
| EB05 | F | 29 | 80 | Birth | Microphthalmia | SC05 | F | 26 | 50 |
| EB06 | M | 28 | 90 | Birth | Congenital Toxoplasmosis | SC06 | M | 29 | 100 |
| EB07 | F | 30 | 60 | Birth | Agenesis | SC07 | F | 27 | 60 |
| EB08 | M | 32 | 80 | Birth | Leber Congenital Amaurosis | SC08 | M | 34 | 70 |
| EB09 | M | 34 | 80 | Birth | Anophthalmia | SC09 | M | 31 | 100 |
| EB10 | F | 38 | 40 | Birth | Retinopathy of Prematurity | SC10 | F | 41 | 60 |
| EB11 | F | 28 | 70 | Birth | Retinopathy of Prematurity | SC11 | F | 26 | 90 |
| EB12 | M | 30 | 70 | Birth | Leber Congenital Amaurosis | SC12 | M | 32 | 80 |
| EB13 | F | 34 | 100 | Birth | Retinopathy of Prematurity | SC13 | F | 31 | 90 |
| EB14 | F | 32 | 100 | 2 Years | Retinoblastoma | SC14 | F | 31 | 70 |
| EB15 | F | 35 | 60 | 4 Years | Retinitis Pigmentosa | SC15 | F | 37 | 100 |
| EB16 | F | 32 | 60 | Birth | Retinitis Pigmentosa | SC16 | F | 33 | 100 |

[a] Handedness was assessed with a modified version of the Edinburgh handedness questionnaire, which is also suitable for the blind population.

**Supplementary Table 2. Stimuli**

| Words | English Translation |
|---|---|
| anello | ring |
| braccialetto | bracelet |
| candela | candle |
| ciotola | bowl |
| coperta | blanket |
| cucchiaio | spoon |
| cuscino | pillow |
| diario | notebook |
| fiammifero | matchstick |
| forchetta | fork |
| gessetto | chalk |
| lavagna | blackboard |
| moneta | coin |
| penna | pen |
| piatto | plate |
| portafoglio | wallet |
| portapenne | penholder |
| salvagente | lifebuoy |
| stuzzicadenti | toothpick |
| timone | rudder |
| tovagliolo | napkin |

**Supplementary Table 3. Inter-rater reliability of object properties and touch experience within each group of participants**

| Rating Items | EB (N = 16) | SC (N = 16) | IS (N = 16) |
|---|---|---|---|
| **Shape Similarity** | ICC = 0.953<br>CI = [0.943, 0.962] | ICC = 0.973<br>CI = [0.968, 0.978] | ICC = 0.973<br>CI = [0.968, 0.978] |
| **Conceptual Association** | ICC = 0.984<br>CI = [0.980, 0.987] | ICC = 0.985<br>CI = [0.982, 0.988] | ICC = 0.985<br>CI = [0.982, 0.988] |
| **Object Size** | ICC = 0.979<br>CI = [0.964, 0.990] | ICC = 0.994<br>CI = [0.989, 0.997] | ICC = 0.992<br>CI = [0.986, 0.996] |
| **Contextual Association** | ICC = 0.613<br>CI = [0.321, 0.817] | ICC = 0.856<br>CI = [0.747, 0.932] | ICC = 0.919<br>CI = [0.859, 0.962] |
| **Toolness** | ICC = 0.911<br>CI = [0.844, 0.958] | ICC = 0.893<br>CI = [0.812, 0.949] | ICC = 0.928<br>CI = [0.874, 0.966] |
| **Touch Experience** | ICC = 0.970<br>CI = [0.948, 0.986] | ICC = 0.975<br>CI = [0.956, 0.988] | ICC = 0.965<br>CI = [0.939, 0.984] |

ICC: the intraclass correlation based on a mean-rating, consistency, two-way random model (i.e., ICC(C,k); McGraw & Wong, 1996)

CI: 95% confidence interval of the true ICC

According to Koo & Li (2016), Poor reliability: < 0.5; Moderate reliability: 0.5-0.75; Good reliability: 0.75-0.9; Excellent reliability: > 0.9

**Supplementary Table 4. Neural representation in bilateral aIPS**

| Three-way Mixed ANOVA * | Left aIPS | | Right aIPS | |
|---|---|---|---|---|
| **Groups** (EB vs. SC) | F(1, 30) = 2.471 | p = 0.126 | F(1, 30) = 2.004 | p = 0.167 |
| **Tasks** (Shape vs. Conceptual) | F(1, 30) = 1.552 | p = 0.223 | F(1, 30) = 0.442 | p = 0.551 |
| **Representations** (Shape vs. Conceptual) | **F(1, 30) = 26.995** | **p < 0.001** | **F(1, 30) = 5.524** | **p = 0.026** |
| **Groups × Tasks** | F(1, 30) = 2.996 | p = 0.094 | F(1, 30) = 0.819 | p = 0.373 |
| **Groups × Representations** | F(1, 30) = 0.021 | p = 0.885 | F(1, 30) = 0.117 | p = 0.735 |
| **Tasks × Representations** | **F(1, 30) = 18.300** | **p < 0.001** | **F(1, 30) = 16.807** | **p < 0.001** |
| **Groups × Tasks × Representations** | F(1, 30) = 1.495 | p = 0.231 | F(1, 30) = 0.367 | p = 0.549 |

* The Groups factor was between-subject, whereas Tasks and Representations were within-subject factors.

**Supplementary Table 5. Neural representation in bilateral pIPS**

| **Three-way Mixed ANOVA** * | **Left pIPS** | | **Right pIPS** | |
|---|---|---|---|---|
| **Groups**<br>(EB vs. SC) | F(1, 30) = 2.925e-5 | p = 0.996 | F(1, 30) = 0.350 | p = 0.559 |
| **Tasks**<br>(Shape vs. Conceptual) | F(1, 30) = 0.840 | p = 0.367 | F(1, 30) = 0.517 | p = 0.478 |
| **Representations**<br>(Shape vs. Conceptual) | **F(1, 30) = 11.158** | **p = 0.002** | **F(1, 30) = 12.745** | **p = 0.001** |
| **Groups × Tasks** | F(1, 30) = 2.203 | p = 0.148 | F(1, 30) = 0.676 | p = 0.417 |
| **Groups × Representations** | F(1, 30) = 1.104 | p = 0.302 | F(1, 30) = 2.087 | p = 0.159 |
| **Tasks × Representations** | **F(1, 30) = 26.524** | **p < 0.001** | **F(1, 30) = 15.401** | **p < 0.001** |
| **Groups × Tasks × Representations** | F(1, 30) = 1.819 | p = 0.188 | **F(1, 30) = 4.597** | **p = 0.040** |

* The Groups factor was between-subject, whereas Tasks and Representations were within-subject factors.

**Supplementary Table 6. Neural representation in bilateral vPMC**

| Three-way Mixed ANOVA * | Left vPMC | | Right vPMC | |
|---|---|---|---|---|
| **Groups**<br>(EB vs. SC) | F(1, 30) = 0.548 | p = 0.465 | F(1, 30) = 0.033 | p = 0.858 |
| **Tasks**<br>(Shape vs. Conceptual) | F(1, 30) = 0.476 | p = 0.496 | F(1, 30) = 0.882 | p = 0.355 |
| **Representations**<br>(Shape vs. Conceptual) | **F(1, 30) = 7.494** | **p = 0.010** | **F(1, 30) = 8.336** | **p = 0.007** |
| **Groups × Tasks** | F(1, 30) = 1.086 | p = 0.306 | F(1, 30) = 3.026 | p = 0.092 |
| **Groups × Representations** | F(1, 30) = 0.197 | p = 0.660 | F(1, 30) = 0.171 | p = 0.682 |
| **Tasks × Representations** | **F(1, 30) = 11.741** | **p = 0.002** | **F(1, 30) = 16.044** | **p < 0.001** |
| **Groups × Tasks × Representations** | F(1, 30) = 2.071 | p = 0.160 | F(1, 30) = 4.102 | p = 0.052 |

* The Groups factor was between-subject, whereas Tasks and Representations were within-subject factors.

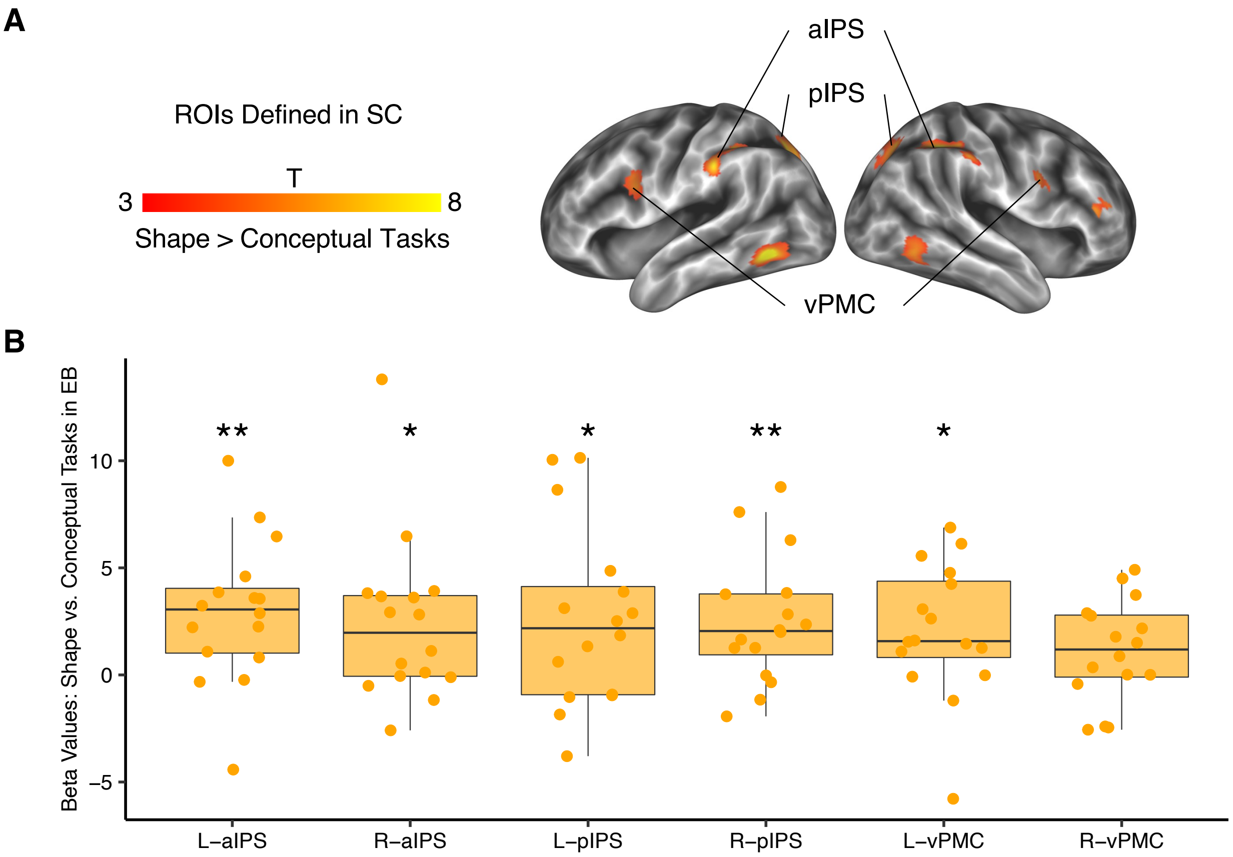


**Supplementary Figure 1.** The vPMC-IPS circuit showed greater activation in the shape task than in the conceptual task in EB. (A) ROIs defined in the contrast between shape and conceptual tasks in SC (vertex-wise $p < 0.001$, cluster-level FWE corrected $p < 0.05$). (B) ROI analyses in the contrast between shape and conceptual tasks in EB using the ROIs defined in SC. *: $p < 0.05$, **: $p < 0.01$.

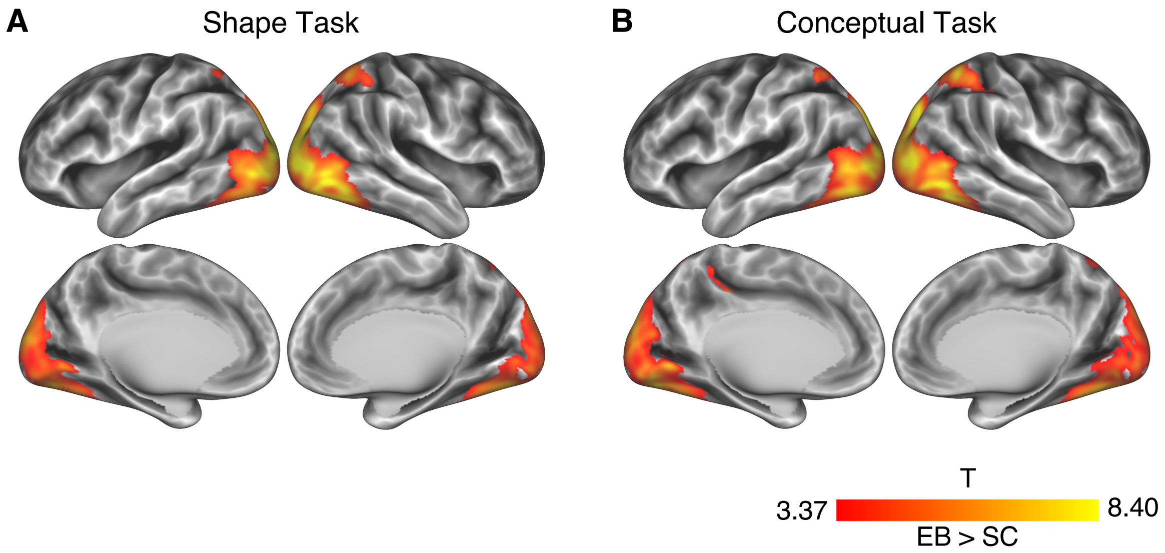


**Supplementary Figure 2.** EB versus SC in shape and conceptual tasks (vertex-wise $p < 0.001$, cluster-level FWE corrected $p < 0.05$). (A) EB versus SC in the shape task. (B) EB versus SC in the conceptual task.

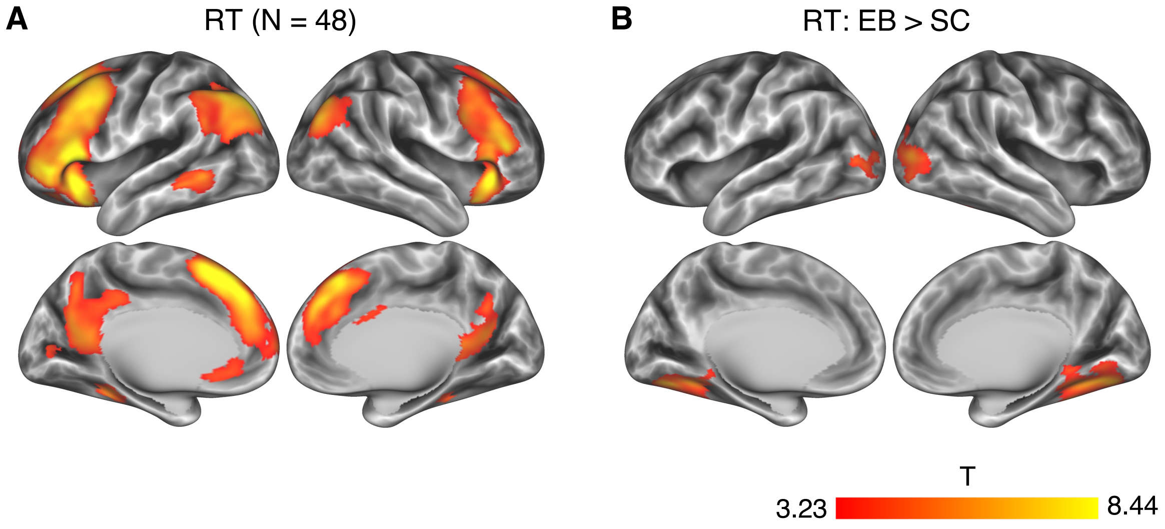


**Supplementary Figure 3.** Neural correlates of reaction time (RT, vertex-wise $p < 0.001$, cluster-level FWE corrected $p < 0.05$). (A) The RT effect across all the participants (N = 48). (B) The differences in the RT effect between the two groups (EB versus SC).

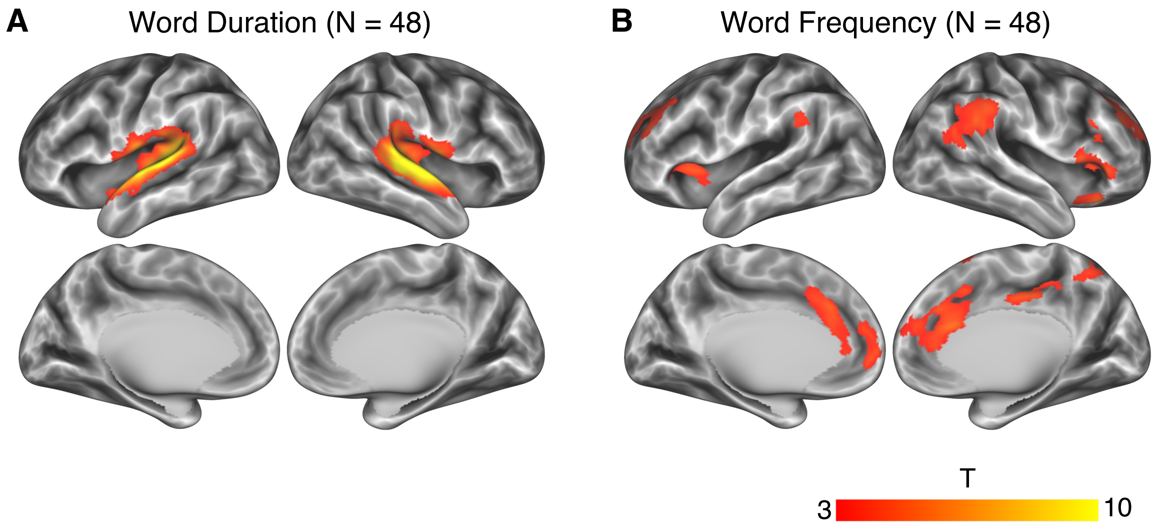


**Supplementary Figure 4.** Neural correlates of linguistic variables (vertex-wise $p < 0.001$, cluster-level FWE corrected $p < 0.05$). (A) Neural correlates of word duration. (B) Neural correlates of word frequency.

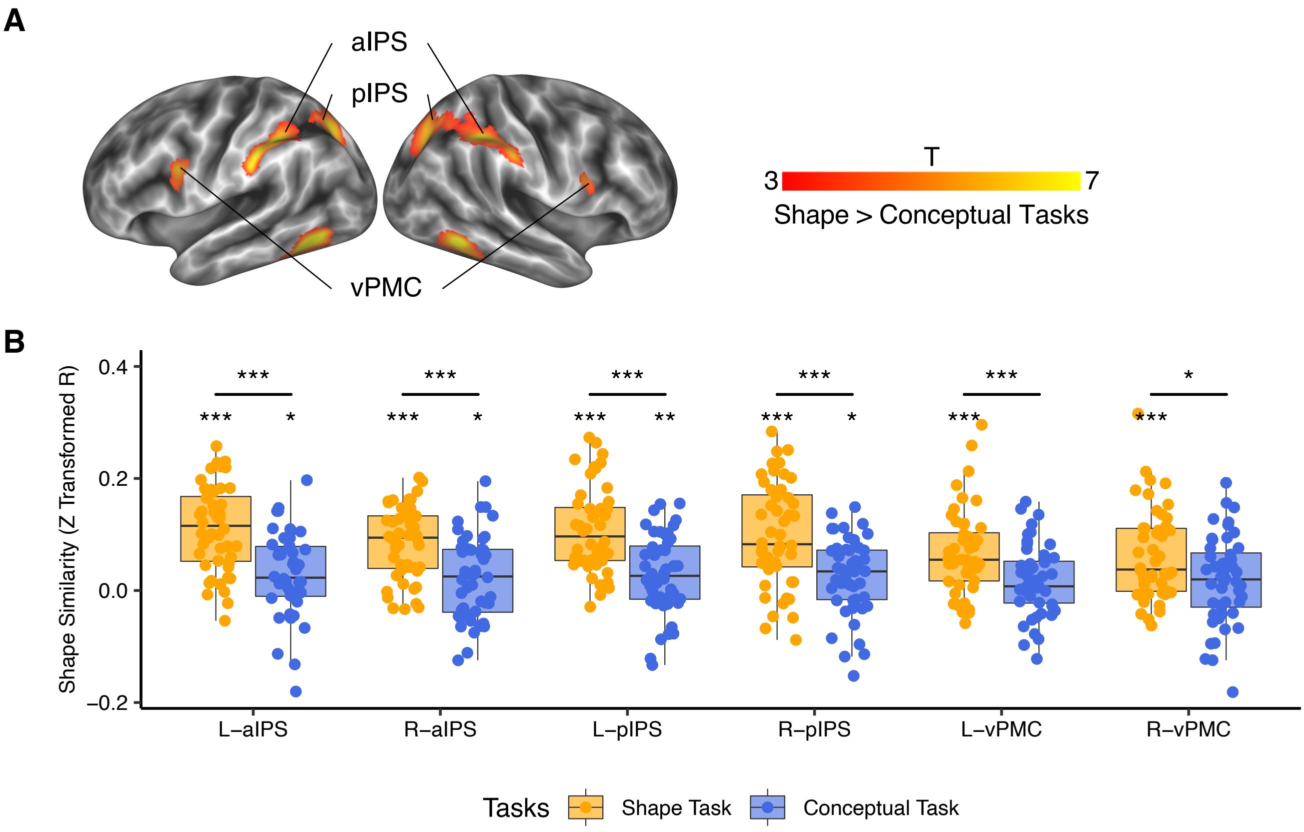


**Supplementary Figure 5.** RSA results of shape similarity in the brain areas with greater activation in the shape task than in the conceptual task. (A) Brain areas with significantly greater activation in the shape task than in the conceptual task defined in Figure 2A. (B) RSA results of these shape-relevant areas in shape and conceptual tasks. *: $p < 0.05$, **: $p < 0.01$, ***: $p <= 0.001$.

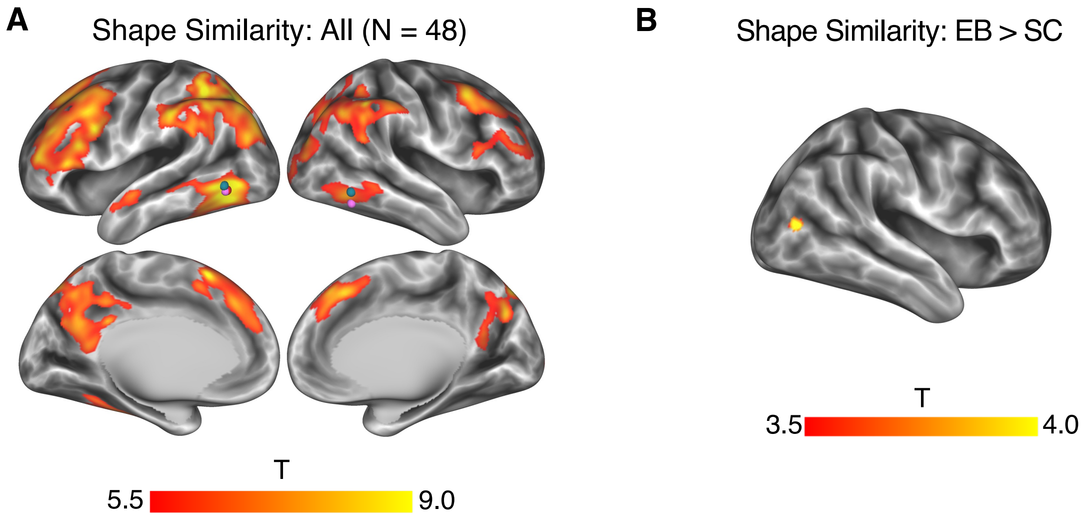


**Supplementary Figure 6.** Whole-brain searchlight results of shape similarity. A. Whole-brain searchlight results of shape similarity across all participants (N = 48; vertex-wise FWE corrected $p < 0.005$, cluster size > 400 $mm^2$). B. Group difference of whole-brain searchlight of shape similarity between EB and SC (vertex-wise $p < 0.001$, cluster-level FWE corrected $p < 0.05$).

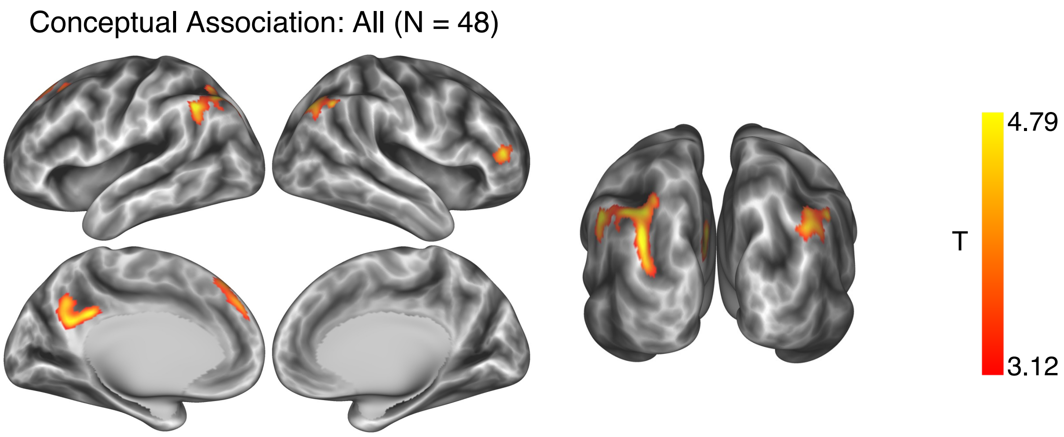


**Supplementary Figure 7.** Whole-brain searchlight results of conceptual association across all participants ($N = 48$; vertex-wise $p < 0.001$, cluster-level FWE corrected $p < 0.05$).

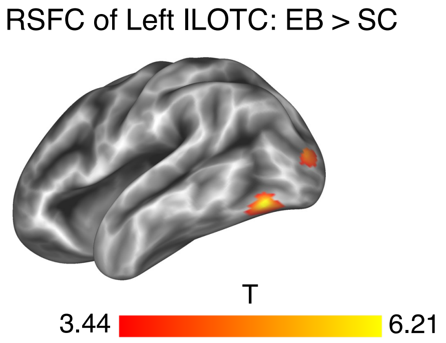


**Supplementary Figure 8.** Contrast between EB and SC in the seed-based RSFC results from the left ILOTC (vertex-wise $p < 0.001$, cluster-level FWE corrected $p < 0.05$).

### Supplementary Text 1. English translation of survey questions

*Shape Familiarity*
To what degree do you know the object's typical shape? (7-point Likert scale rating)
1: do not know it at all 7: know it very well

*Conceptual Familiarity*
To which degree do you know what this object is used for? (7-point Likert scale rating)
1: do not know it at all 7: know it very well

*Touch Experience*
How frequently have you touched this object? (7-point Likert scale rating)
1: have never touched it before 7: touch it every day

*Size*
How big is this object? (7-point Likert scale rating)
1: as small as a needle 7: as big as a television

*Contextual Association*
To which extent is this object associated with a specific context? (7-point Likert scale rating)
For example, a "cellphone" can occur in many different contexts and is unassociated with any particular cortex. You might rate 1 or 2. Instead, a "bowling ball" can only occur on the bowling alley; it is strongly associated with one specific context. You might rate 7.

*Toolness*
To which extent is this object a tool? (7-point Likert scale rating)
A tool is defined as a graspable and manipulable object that can transform the motor output into predictable mechanical actions for the purposes of attaining specific goals. You might rate 1 for "lamp," "chair," and "clock"; 7 for "hammer," "saw," and "drill."

*Pairwise Shape Similarity*
To which extent is this pair of objects similar in shape? (7-point Likert scale rating)
For example, "racket" and "pan" are similar in shape; you might rate 6 or 7; "racket" and "tennis ball" are not similar in shape, you might rate 1 or 2. Due to the recent expansion of business innovation, some objects may exist in a variety of shapes, e.g., watermelon may be in a square shape. Please rate the shape based on its most typical shape (i.e., spherical watermelon). Please disregard the other object properties, e.g., color, size, texture, and function.

*Pairwise Conceptual Association*
To which extent is this pair of objects conceptual associated? (7-point Likert scale rating)
For example, "racket" and "tennis ball" are conceptually associated; you might rate 7; "racket" and "pan" are not conceptually associated, you might rate 1. Please disregard the other object properties, e.g., color, size, texture, and shape.

### Supplementary Text 2. MRI preprocessing using fMRIPrep

Results included in this manuscript come from preprocessing performed using fMRIPrep 20.0.5 (Esteban, Markiewicz, et al. (2018); Esteban, Blair, et al. (2018); RRID:SCR_016216), which is based on Nipype 1.4.2 (Gorgolewski et al. (2011); Gorgolewski et al. (2018); RRID:SCR_002502).

*Anatomical data preprocessing*

The T1-weighted (T1w) image was corrected for intensity non-uniformity (INU) with N4BiasFieldCorrection (Tustison et al. 2010), distributed with ANTs 2.2.0 (Avants et al. 2008, RRID:SCR_004757), and used as T1w-reference throughout the workflow. The T1w-reference was then skull-stripped with a Nipype implementation of the antsBrainExtraction.sh workflow (from ANTs), using OASIS30ANTs as target template. Brain tissue segmentation of cerebrospinal fluid (CSF), white-matter (WM) and gray-matter (GM) was performed on the brain-extracted T1w using fast (FSL 5.0.9, RRID:SCR_002823, Zhang, Brady, and Smith 2001). Brain surfaces were reconstructed using recon-all (FreeSurfer 6.0.1, RRID:SCR_001847, Dale, Fischl, and Sereno 1999), and the brain mask estimated previously was refined with a custom variation of the method to reconcile ANTs-derived and FreeSurfer-derived segmentations of the cortical gray-matter of Mindboggle (RRID:SCR_002438, Klein et al. 2017). Volume-based spatial normalization to two standard spaces (MNI152NLin6Asym, MNI152NLin2009cAsym) was performed through nonlinear registration with antsRegistration (ANTs 2.2.0), using brain-extracted versions of both T1w reference and the T1w template. The following templates were selected for spatial normalization: FSL's MNI ICBM 152 non-linear 6th Generation Asymmetric Average Brain Stereotaxic Registration Model [Evans et al. (2012), RRID:SCR_002823; TemplateFlow ID: MNI152NLin6Asym], ICBM 152 Nonlinear Asymmetrical template version 2009c [Fonov et al. (2009), RRID:SCR_008796; TemplateFlow ID: MNI152NLin2009cAsym],

*Functional data preprocessing*

For each of the 11 BOLD runs found per subject (across all tasks and sessions), the following preprocessing was performed. First, a reference volume and its skull-stripped version were generated using a custom methodology of fMRIPrep. A B0-nonuniformity map (or fieldmap) was estimated based on two (or more) echo-planar imaging (EPI) references with opposing phase-encoding directions, with 3dQwarp Cox and Hyde (1997) (AFNI 20160207). Based on the estimated susceptibility distortion, a corrected EPI (echo-planar imaging) reference was calculated for a more accurate co-registration with the anatomical reference. The BOLD reference was then co-registered to the T1w reference using bbregister (FreeSurfer) which implements boundary-based registration (Greve and Fischl 2009). Co-registration was configured with six degrees of freedom. Head-motion parameters with respect to the BOLD reference (transformation matrices, and six corresponding rotation and translation parameters) are estimated before any spatiotemporal filtering using mcflirt (FSL 5.0.9, Jenkinson et al. 2002). The BOLD time-series were resampled onto the following surfaces (FreeSurfer reconstruction nomenclature): fsaverage5, fsnative, fsaverage. The BOLD time-series (including slice-timing correction when applied) were resampled onto their original, native space by applying a single, composite transform to correct for head-motion and susceptibility distortions. These resampled BOLD time-series will be referred to as preprocessed BOLD in original space, or just preprocessed BOLD. Grayordinates files (Glasser et al. 2013) containing 91k samples were also generated using the highest-resolution fsaverage as intermediate standardized surface space. Several confounding time-series were calculated based on the preprocessed BOLD: framewise displacement (FD), DVARS and three region-wise global signals. FD and DVARS are calculated for each functional run, both using their implementations in Nipype (following the definitions by Power et al. 2014). The three global signals are extracted within the CSF, the WM, and the whole-brain masks. Additionally, a set of physiological regressors were extracted to allow for component-based noise correction (CompCor, Behzadi et al. 2007). Principal components are estimated after high-pass filtering the preprocessed BOLD time-series (using a discrete cosine filter with 128s cut-off) for the two CompCor variants: temporal (tCompCor) and anatomical (aCompCor). tCompCor components are then calculated from the top 5% variable voxels within a mask covering the subcortical regions.

This subcortical mask is obtained by heavily eroding the brain mask, which ensures it does not include cortical GM regions. For aCompCor, components are calculated within the intersection of the aforementioned mask and the union of CSF and WM masks calculated in T1w space, after their projection to the native space of each functional run (using the inverse BOLD-to-T1w transformation). Components are also calculated separately within the WM and CSF masks. For each CompCor decomposition, the k components with the largest singular values are retained, such that the retained components' time series are sufficient to explain 50 percent of variance across the nuisance mask (CSF, WM, combined, or temporal). The remaining components are dropped from consideration. The head-motion estimates calculated in the correction step were also placed within the corresponding confounds file. The confound time series derived from head motion estimates and global signals were expanded with the inclusion of temporal derivatives and quadratic terms for each (Satterthwaite et al. 2013). Frames that exceeded a threshold of 0.5 mm FD or 1.5 standardised DVARS were annotated as motion outliers. All resamplings can be performed with a single interpolation step by composing all the pertinent transformations (i.e. head-motion transform matrices, susceptibility distortion correction when available, and co-registrations to anatomical and output spaces). Gridded (volumetric) resamplings were performed using antsApplyTransforms (ANTs), configured with Lanczos interpolation to minimize the smoothing effects of other kernels (Lanczos 1964). Non-gridded (surface) resamplings were performed using mri_vol2surf (FreeSurfer).

Many internal operations of fMRIPrep use Nilearn 0.6.2 (Abraham et al. 2014, RRID:SCR_001362), mostly within the functional processing workflow. For more details of the pipeline, see the section corresponding to workflows in fMRIPrep's documentation.

*Copyright Waiver*

The above boilerplate text was automatically generated by fMRIPrep with the express intention that users should copy and paste this text into their manuscripts unchanged. It is released under the CC0 license.

*References*

Abraham, Alexandre, Fabian Pedregosa, Michael Eickenberg, Philippe Gervais, Andreas Mueller, Jean Kossaifi, Alexandre Gramfort, Bertrand Thirion, and Gael Varoquaux. 2014. "Machine Learning for Neuroimaging with Scikit-Learn." Frontiers in Neuroinformatics 8. https://doi.org/10.3389/fninf.2014.00014.

Avants, B.B., C.L. Epstein, M. Grossman, and J.C. Gee. 2008. "Symmetric Diffeomorphic Image Registration with Cross-Correlation: Evaluating Automated Labeling of Elderly and Neurodegenerative Brain." Medical Image Analysis 12 (1): 26–41. https://doi.org/10.1016/j.media.2007.06.004.

Behzadi, Yashar, Khaled Restom, Joy Liau, and Thomas T. Liu. 2007. "A Component Based Noise Correction Method (CompCor) for BOLD and Perfusion Based fMRI." NeuroImage 37 (1): 90–101. https://doi.org/10.1016/j.neuroimage.2007.04.042.

Cox, Robert W., and James S. Hyde. 1997. "Software Tools for Analysis and Visualization of fMRI Data." NMR in Biomedicine 10 (4-5): 171–78. https://doi.org/10.1002/(SICI)1099-1492(199706/08)10:4/5<171::AID-NBM453>3.0.CO;2-L.

Dale, Anders M., Bruce Fischl, and Martin I. Sereno. 1999. "Cortical Surface-Based Analysis: I. Segmentation and Surface Reconstruction." NeuroImage 9 (2): 179–94. https://doi.org/10.1006/nimg.1998.0395.

Esteban, Oscar, Ross Blair, Christopher J. Markiewicz, Shoshana L. Berleant, Craig Moodie, Feilong Ma, Ayse Ilkay Isik, et al. 2018. "FMRIPrep." Software. Zenodo. https://doi.org/10.5281/zenodo.852659.

Esteban, Oscar, Christopher Markiewicz, Ross W Blair, Craig Moodie, Ayse Ilkay Isik, Asier Erramuzpe Aliaga, James Kent, et al. 2018. "fMRIPrep: A Robust Preprocessing Pipeline for Functional MRI." Nature Methods. https://doi.org/10.1038/s41592-018-0235-4.

Evans, AC, AL Janke, DL Collins, and S Baillet. 2012. "Brain Templates and Atlases." NeuroImage 62 (2): 911–22. https://doi.org/10.1016/j.neuroimage.2012.01.024.

Fonov, VS, AC Evans, RC McKinstry, CR Almli, and DL Collins. 2009. “Unbiased Nonlinear Average Age-Appropriate Brain Templates from Birth to Adulthood.” NeuroImage 47, Supplement 1: S102. https://doi.org/10.1016/S1053-8119(09)70884-5.

Glasser, Matthew F., Stamatios N. Sotiropoulos, J. Anthony Wilson, Timothy S. Coalson, Bruce Fischl, Jesper L. Andersson, Junqian Xu, et al. 2013. “The Minimal Preprocessing Pipelines for the Human Connectome Project.” NeuroImage, Mapping the connectome, 80: 105–24. https://doi.org/10.1016/j.neuroimage.2013.04.127.

Gorgolewski, K., C. D. Burns, C. Madison, D. Clark, Y. O. Halchenko, M. L. Waskom, and S. Ghosh. 2011. “Nipype: A Flexible, Lightweight and Extensible Neuroimaging Data Processing Framework in Python.” Frontiers in Neuroinformatics 5: 13. https://doi.org/10.3389/fninf.2011.00013.

Gorgolewski, Krzysztof J., Oscar Esteban, Christopher J. Markiewicz, Erik Ziegler, David Gage Ellis, Michael Philipp Notter, Dorota Jarecka, et al. 2018. “Nipype.” Software. Zenodo. https://doi.org/10.5281/zenodo.596855.

Greve, Douglas N, and Bruce Fischl. 2009. “Accurate and Robust Brain Image Alignment Using Boundary-Based Registration.” NeuroImage 48 (1): 63–72. https://doi.org/10.1016/j.neuroimage.2009.06.060.

Jenkinson, Mark, Peter Bannister, Michael Brady, and Stephen Smith. 2002. “Improved Optimization for the Robust and Accurate Linear Registration and Motion Correction of Brain Images.” NeuroImage 17 (2): 825–41. https://doi.org/10.1006/nimg.2002.1132.

Klein, Arno, Satrajit S. Ghosh, Forrest S. Bao, Joachim Giard, Yrjö Häme, Eliezer Stavsky, Noah Lee, et al. 2017. “Mindboggling Morphometry of Human Brains.” PLOS Computational Biology 13 (2): e1005350. https://doi.org/10.1371/journal.pcbi.1005350.

Lanczos, C. 1964. “Evaluation of Noisy Data.” Journal of the Society for Industrial and Applied Mathematics Series B Numerical Analysis 1 (1): 76–85. https://doi.org/10.1137/0701007.

Power, Jonathan D., Anish Mitra, Timothy O. Laumann, Abraham Z. Snyder, Bradley L. Schlaggar, and Steven E. Petersen. 2014. “Methods to Detect, Characterize, and Remove Motion Artifact in Resting State fMRI.” NeuroImage 84 (Supplement C): 320–41. https://doi.org/10.1016/j.neuroimage.2013.08.048.

Satterthwaite, Theodore D., Mark A. Elliott, Raphael T. Gerraty, Kosha Ruparel, James Loughead, Monica E. Calkins, Simon B. Eickhoff, et al. 2013. “An improved framework for confound regression and filtering for control of motion artifact in the preprocessing of resting-state functional connectivity data.” NeuroImage 64 (1): 240–56. https://doi.org/10.1016/j.neuroimage.2012.08.052.

Tustison, N. J., B. B. Avants, P. A. Cook, Y. Zheng, A. Egan, P. A. Yushkevich, and J. C. Gee. 2010. “N4ITK: Improved N3 Bias Correction.” IEEE Transactions on Medical Imaging 29 (6): 1310–20. https://doi.org/10.1109/TMI.2010.2046908.

Zhang, Y., M. Brady, and S. Smith. 2001. “Segmentation of Brain MR Images Through a Hidden Markov Random Field Model and the Expectation-Maximization Algorithm.” IEEE Transactions on Medical Imaging 20 (1): 45–57. https://doi.org/10.1109/42.906424.
